# 3-hydroxyanthranilic acid – a new metabolite for healthy lifespan extension

**DOI:** 10.1101/2021.06.01.446651

**Authors:** Hope Dang, Raul Castro-Portuguez, Luis Espejo, Grant Backer, Sam Freitas, Erica Spence, Jeremy Meyers, Karissa Shuck, Caroline Corban, Teresa Liu, Shannon Bean, Susan Sheehan, Ron Korstanje, George L. Sutphin

## Abstract

The metabolism of tryptophan to nicotinamide adenine dinucleotide (NAD^+^) through the kynurenine pathway is increasingly linked to aging and age-associated disease. Kynurenine pathway enzymes and metabolites influence a range of molecular processes critical to healthy aging, including regulation of inflammatory and immune responses, cellular redox homeostasis, and energy production. Aberrant kynurenine metabolism is observed during normal aging and has been implicated in a range of age-associated pathologies, including chronic inflammation, atherosclerosis, neurodegeneration, and cancer. In previous work, we and others identified three genes—*kynu-1, tdo-2*, and *acsd-1*—encoding kynurenine pathway enzymes for which decreasing expression extends lifespan in invertebrate models. Here we report that knockdown of *haao-1*, a fourth kynurenine pathway gene encoding the enzyme 3-hydroxyanthranilic acid dioxygenase (HAAO), extends lifespan by ~30% and delays age-associated decline in health in *Caenorhabditis elegans*. This lifespan extension is mediated by increased physiological levels of the HAAO substrate 3-hydroxyanthranilic acid (3HAA). 3HAA increases resistance to oxidative stress during aging by directly degrading hydrogen peroxide and activating the Nrf2/SKN-1 oxidative stress response. Aging mice fed a diet supplemented with 3HAA are similarly long-lived. Our results identify HAAO and 3HAA as novel therapeutic targets for age-associated disease. This works provides a foundation for more detailed examination of the molecular mechanisms underlying the benefits of 3HAA, and how these mechanisms interact with other interventions both within and beyond the kynurenine pathway. We anticipate that these findings will bolster growing interest in developing pharmacological strategies to target tryptophan metabolism to improve health aging.

## Introduction

Physiological tryptophan (TRP) has a variety of potential fates in eukaryotic cells, including incorporation into proteins, conversion to serotonin or melatonin, conversion to tryptamine, or conversion to nicotinamide adenine dinucleotide (NAD^+^) through the kynurenine pathway (**Fig. 1A**). The majority (>90%) of ingested tryptophan is catabolized by the kynurenine pathway in mammals^1^. Altered kynurenine pathway activity has been implicated in a range of age-associated diseases in humans, including cardiovascular disease, kidney disease, cancer, and neurodegeneration^2,3^. The structure of the kynurenine pathway is evolutionarily conserved from bacteria through mammals, with a few notable exceptions. The initial, rate-limiting conversion of TRP to N-formylkynurenine (NFK) is carried out by the enzyme tryptophan 2,3-dioxygenase (TDO2) across species. Vertebrate genomes encode a second cytokine-responsive enzyme, IDO1, that catalyzes this reaction, and vertebrate kynurenine pathway initiation is largely segregated between immune cells (IDO1) and liver (TDO2)^4^. A duplication event in the mammalian lineage produced IDO2, which is more ubiquitously expressed and less active than IDO1 and does not respond to cytokines. IDO2’s function is currently less well-defined^4^. Arylformamidase (AFMID) catabolizes NFK to kynurenine (KYN), which is in turn processed to either kynurenic acid (KA) or NAD^+^ through the two major branches of the kynurenine pathway. The NAD^+^ branch converts KYN to NAD^+^ through a series of metabolic steps catalyzed by the enzymes kynurenine 3-monooxygenase (KMO), kynureninase (KYNU), and 3-hydroxyanthranilic acid dioxygenase (HAAO).

**Figure 1.**
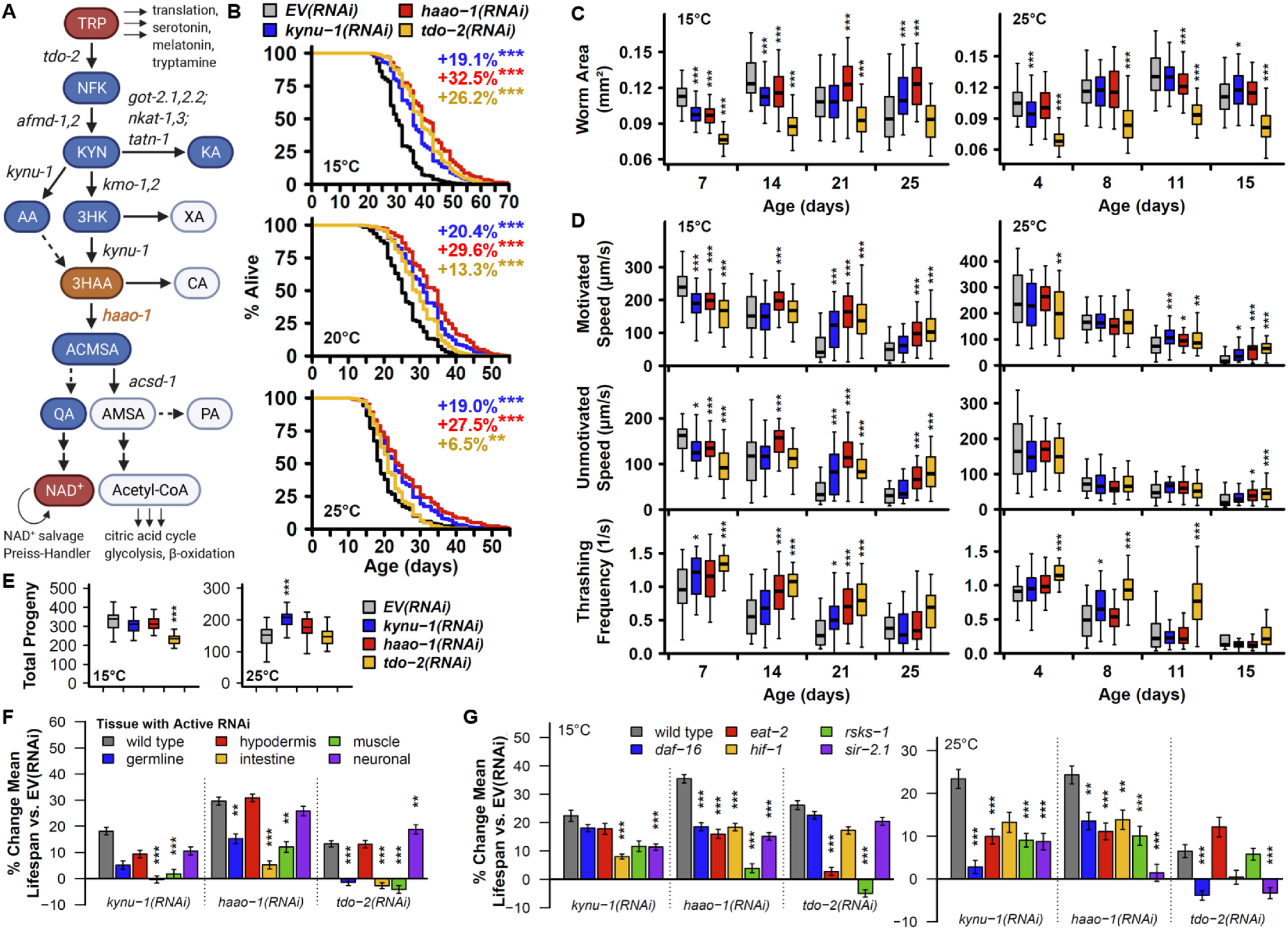
Knockdown of *haao-1* extends healthy lifespan in *C. elegans*. (**A**) Kynurenine pathway schematic (created BioRender). Metabolites are shown in bubbles. *C. elegans* gene names for enzymatic steps are shown in italic text. RNAi knockdown of *kynu-1, haao-1*, or *tdo-2*: (**B**) extends lifespan at 15°C, 20°C, and 25°C (colored text indicates change in mean lifespan relative to control); (**C**) slows growth but allows worms to maintain body size (i.e. worm area) later in life; (**D**) slows motivated and unmotivated crawl speed early in life but allows worms to maintain speed and thrashing frequency in liquid later in life. (**E**) *tdo-2(RNAi)* reduces brood size at 15°C while *kynu-1(RNAi)* increased brood size at 25°C. Lifespan extension from RNAi knockdown of *kynu-1, haao-1*, or *tdo-2* displays different patterns of dependence on (**F**) tissue and (**G**) genes in established aging pathways. All error bars indicate standard error of mean. * p < 0.05, ** p < 0.01, *** p < 0.001. Abbreviations: TRP = tryptophan, NFK = N-formylkynurenine, KYN = kynurenine, KA = kynurenic acid, AA = anthranilic acid, 3HK = 3-hydroxykynurenine, XA = xanthurenic acid, 3HAA = 3-hydroxyanthranilic acid, CA = cinnabarinic acid, ACMSA = aminocarboxymuconate semialdehyde, AMSA = aminomuconic semialdehyde, QA = quinolinic acid, Acetyl-CoA = acetyl coenzyme A, NAD^+^ = nicotinamide adenine dinucleotide.

Several kynurenine pathway interventions provide direct benefits in aging invertebrate models. Oxenkrug^5^ observed that common *Drosophila melanogaster* strains with altered eye color resulting from mutations in the gene *vermillion* (*ν*), encoding the fruit fly ortholog of TDO2, were long-lived. They later reported lifespan extension in wild type *Drosophila* treated with the TDO2 inhibitors α-methyltryptophan (αMT) or 5-methyltryptophan (5MT)^6^. van der Goot et al.^7^ similarly found that RNAi knockdown of the *C. elegans ν* ortholog, *tdo-2*, extended lifespan and improved α-synuclein pathology, likely mediated by elevated TRP^7^. We later discovered that knockdown of *kynu-1*, encoding the enzyme kynureninase (KYNU), can extend healthy *C. elegans* lifespan^8^. Finally, Katsyuba et al.^9^ recently reported lifespan extension following knockdown of *acsd-1*, encoding aminocarboxymuconate semialdehyde decarboxylase (ACMSD), which controls one branchpoint of the kynurenine pathway, catabolizing aminocarboxymuconate semialdehyde (ACMSA) to aminomuconic semialdehyde (AMSA) and subsequently into picolinic acid or acetyl coenzyme-A (**Fig. 1A**). ACMSA is unstable and can also non-enzymatically convert to quinolinic acid (QA), which feeds into the Preiss-Handler NAD^+^ biosynthetic pathway. As a consequence, reduced ACMSD activity results in elevated NAD^+^ production, the proposed mediator of lifespan extension^9^. Knockdown of *acsd-1* has the opposite effect on physiological NAD^+^ levels compared to knockdown of either *tdo-2* or *kynu-1*, implying that kynurenine metabolism can influence aging through both NAD^+^-dependent and NAD^+^-independent mechanisms.

Here we report that a fourth kynurenine pathway enzyme, HAAO, can influence aging in *C. elegans*. Knockout or knockdown of the gene encoding HAAO, *haao-1*, robustly increased lifespan by increasing physiological levels of its substrate metabolite, 3-hydoxyanthranilic acid (3HAA), a mechanism distinct from previous reports of lifespan extension resulting from targeting kynurenine pathway enzymes. Worms with reduced *haao-1* activity maintained biomarkers of health later in life. We further present evidence mechanistically implicating heightened oxidative stress resistance in the beneficial effects of 3HAA. Finally, we find the dietary supplementation with 3HAA extended lifespan in a small cohort of aging mice.

## Results

### Knockdown of *haao-1* extends healthy lifespan in *C. elegans*

In previous work we reported that knockdown of *kynu-1* robustly increased lifespan through mechanisms apparently distinct from knockdown of *tdo-2*^8^. Our initial goal in this work was to examine other intervention targets in the kynurenine pathway with benefits to longevity and age-associated disease. In vertebrate and most invertebrate model systems for aging, three enzymes aside from TDO2 and KYNU are encoded by singe genes and thus amenable to straightforward knockdown using RNAi: AFMID, KMO, and HAAO. However, duplication events in the nematode lineage resulted in both AFMID and KMO being encoded by two genes each in *C. elegans* (**Fig. 1A**)^10,11^. We therefore focused on *haao-1*, encoding the enzyme HAAO in *C. elegans*. RNAi knockdown of *haao-1* robustly extended lifespan by ~30%. This effect was consistently greater than that observed for either *kynu-1(RNAi)* or *tdo-2(RNAi)*, and—like *kynu-1(RNAi)* but unlike *tdo-2(RNAi)*^8^—was consistent across the viable temperature range for *C. elegans* (**Fig. 1B**). We validated both *haao-1(RNAi)* and *kynu-1(RNAi)* by confirming that fluorescence in HAAO-1::wrmscarlet and KYNU-1::wrmscarlet transgenic fusion strains was reduced below detectable levels (**Extended Data Fig. 5B**). At 15°C, *haao-1(RNAi)* animals were slow to reach full body size, but resistant to the age-dependent reduction in body size observed in animals subjected to empty vector RNAi *(EV(RNAi))* (**Fig. 1C**). Similarly, *haao-1(RNAi)* animals were slower than *EV(RNAi)* animals during early adulthood in the context of both motivated (stimulated by dropping the petri plate from a height of ~1 inch) and unmotivated crawling speed on solid media, but resistant to age-dependent reduction in motivated speed, unmotivated speed, and thrashing in liquid media (**Fig. 1D**). These healthspan benefits were blunted at 25°C, resembling the pattern for *kynu-1(RNAi)* but distinct from *tdo-2(RNAi)* (**Fig. 1C-D**). In previous work we observed a reduced brood size in animals subjected to *tdo-2(RNAi)* at 15°C and an increased brood size for animals subjected to *kynu-1(RNAi)*, suggesting an interaction between kynurenine metabolism, environmental temperature, and reproductive capacity. We did not observe a difference in total brood size in response to *haao-1(RNAi)* (**Fig. 1E**); however, the time-course of egg production suggested a slightly delayed egg production profile relative to *EV(RNAi)*, similar to *kynu-1(RNAi)* but less pronounced that *tdo-2(RNAi)* (**Extended Data Fig. 1A**).

To further characterize the relationship between aging and kynurenine metabolism, we measured the impact *kynu-1(RNAi), haao-1(RNAi)*, and *tdo-2(RNAi)* on lifespan is a set of *C. elegans* strains with molecular machinery for RNAi globally inactivated and reactivated in restricted tissues, allowing tissue-specific RNAi knockdown. For all three genes, RNAi knockdown in either the hypodermis or neurons was sufficient to extend lifespan to a similar degree as whole-body knockdown. RNAi knockdown in intestine or muscle was insufficient to extend lifespan for *kynu-1* or *tdo-2*, and produced intermediate lifespan extension for *haao-1* (**Fig. 1F**, **Extended Data Fig. 2**). Germline-specific knockdown generated a similar pattern to intestine and muscle, except that variability between replicates made the outcome indeterminate for *kynu-1(RNAi)*. This pattern suggests that the impact of kynurenine pathway interventions in *C. elegans* is likely driven by activity in hypodermis, neurons, and possibly other tissues not examined in this work.

We next examined the interaction between kynurenine metabolism and genetic pathways with an established role in aging by measuring lifespan in response to *kynu-1(RNAi)*, *haao-1(RNAi)*, and *tdo-2(RNAi)* for strains with the following mutations: *daf-16(mu68)* (impaired insulin/IGF-1-like signaling), *eat-2(ad465)* (a genetic model of dietary restriction with reduced pharyngeal pumping), *hif-1(ia4)* (impaired hypoxic response), *rsks-1(ok1255)* (encoding the ribosomal protein S6 kinase, which regulates translation in response to mTOR signaling), and *sir-2.1(ok434)* (encoding the *C. elegans* ortholog if Sir2). As we previously reported, *kynu-1* and *tdo-2* display distinct and temperature-dependent patterns of interaction with this set of mutations^8^. The pattern of interaction for *haao-1* was distinct from either gene. The lifespan extension resulting from *haao-1(RNAi)* was smaller in all genetic backgrounds tested at both 15°C and 25°C than in wild type worms, and absent in *rsks-1* loss-of-function worms at 15°C and *sir-2.1* worms at 25°C (**Fig. 1G**, **Extended Data Fig. 3, 4**). This pattern of interaction suggests a complex relationship with established aging pathways and does not clearly point to a primary single mediator within the set of pathways examined. Given this data, we decided to approach identifying key mediators of the effect of *haao-1* on aging from the opposite direction, by examining the mechanistic step directly impacted by *haao-1* knockdown.

### Elevated 3HAA mediates lifespan extension from *haao-1* knockdown

Throughout our work with *haao-1*, we noted that animals with reduced *haao-1* activity developed a red coloring that became visible near the head under brightfield illumination on a stereoscope around day 4 of adulthood (data not shown) and accumulated throughout the body with age (**Fig. 2A**, **Extended Data Fig. 5A**). The structure of the kynurenine pathway suggests that genetically inhibiting HAAO activity should result in physiological accumulation of its substrate metabolite, 3HAA (**Fig. 1A**). We purchased purified 3HAA and added it to the solid nematode growth media (NGM) used to culture worms. 3HAA proved to be red in color (**Fig. 2B**), consistent with 3HAA causing the red coloration in worms lacking active *haao-1*. Using liquid chromatography tandem mass-spectrometry (LC-MS/MS) we further confirmed that 3HAA is elevated by ~10-fold in worms subjected to *haao-1(RNAi)*, but not *kynu-1(RNAi)* or *tdo-2(RNAi)* (**Fig. 2C**). We hypothesized that accumulation of 3HAA may mediate lifespan extension from *haao-1* knockdown. In support of this hypothesis, we found that worms maintained on media supplemented with 0.01 to 2.5 mM 3HAA were long-lived relative to untreated animals (**Fig. 2E**, **Extended Data Fig. 5C, D**). Supplementation with 1 mM 3HAA closely mimicked lifespan extension from *haao-1(RNAi)* or the null mutation *haao-1(tm4627)* (**Fig. 2E**). If 3HAA mediates the aging benefits of decreasing HAAO activity, but not KYNU or TDO2, 3HAA supplementation should further increase the already long lifespans of animals with reduced *kynu-1* or *tdo-2*, but not animals with reduced *haao-1*. Our observations confirmed this prediction, with the note that 3HAA slightly extended lifespan in *haao-1* null animals, which we speculate may be due to a suboptimal 3HAA level resulting from absent HAAO activity alone (**Fig. 3F**, **Extended Data Fig. 6**). Supporting this notion and further validating our model that 3HAA mediates the health benefits from *haao-1* knockdown, the full lifespan extension from 3HAA supplementation was rescued in *haao-1* knockout animals when 3HAA production was blocked by the further deletion of *kynu-1* (**Fig. 1A, 3F**, **Extended Data Fig. 6**).

**Figure 2.**
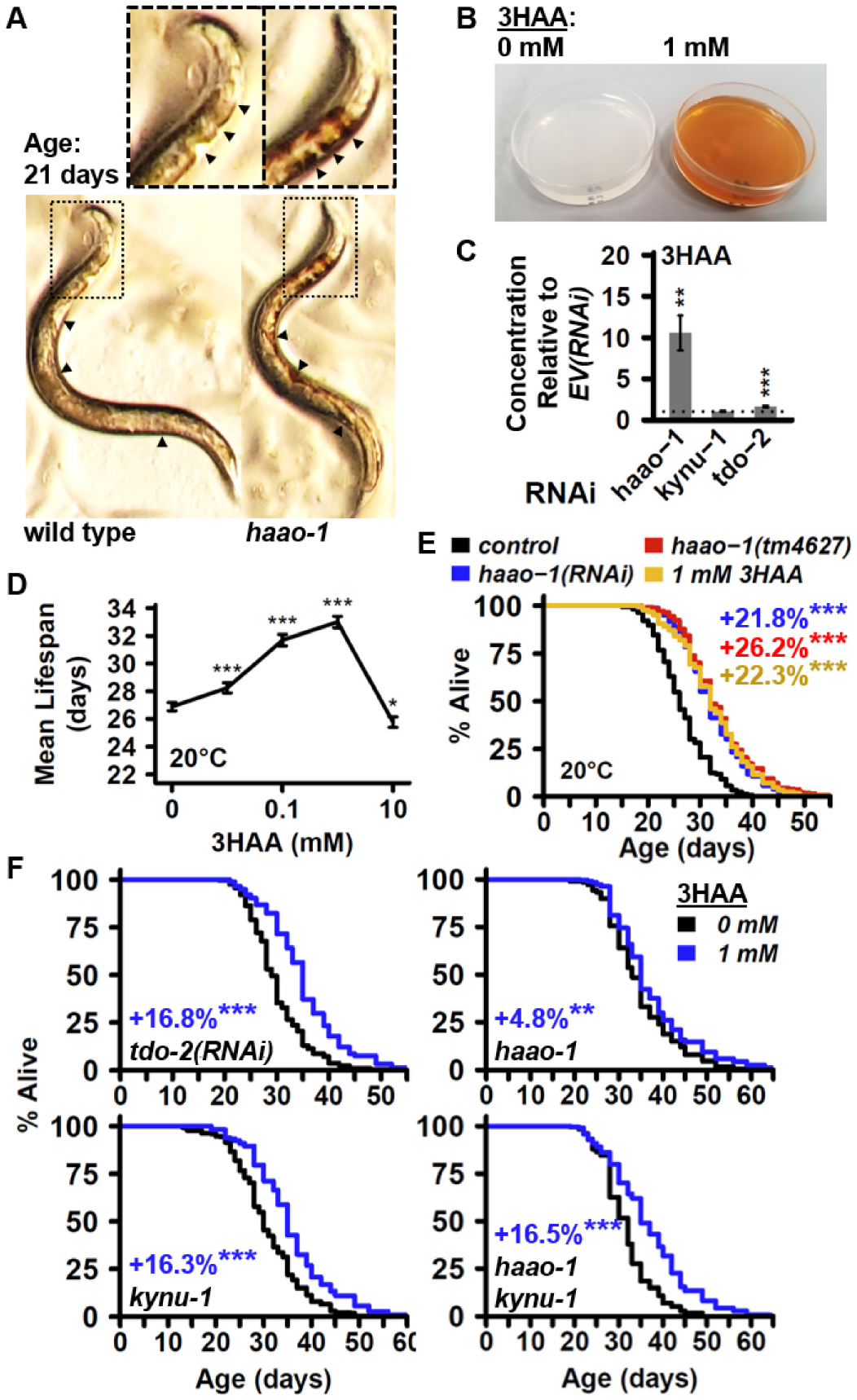
Elevating 3HAA protects against oxidative stress by directly degrading hydrogen peroxide and activating SKN-1-mediated oxidative stress response. (**A**) RNAi knockdown of *haao-1*, deletion of *haao-1 (haao-1(tm4637))*, and media supplemented with 1 mM 3HAA increase survival of *C. elegans* challenged with 20 mM paraquat (colored text indicates change in mean lifespan relative to control). (**B**) 500 μM H_2_O_2_ reduces visible red color of 1 mM 3HAA in water. (**C**) 3HAA reduces detectable H_2_O_2_ in water in a dose-dependent manner. (**D**) Deletion of *haao-1 (haao-1(tm4637))* or 1 mM 3HAA supplementation reduces the age-dependent increase in H_2_O_2_ secretion in *C. elegans*. (**E**) RNAi knockdown of *haao-1* or 1 mM 3HAA supplementation increases fluorescence in *C. elegans* strains transgenic ally expressing the *skn-1::GFP* fusion protein or the *gcs-1p::GFP* or *gst-4p::GFP* promoter activity reporters. All error bars indicate standard error of mean. * p < 0.05, ** p < 0.01, *** p < 0.001.

### 3HAA increases resistance to oxidative stress

We next asked what mechanisms may mediate lifespan extension from 3HAA. Accumulation of reactive oxygen species and dysregulation of cellular oxidative stress response during aging are causally implicated in a variety of age-associated pathologies^12^. 3HAA is a redox active metabolite with either pro- or antioxidant properties depending on physiological context, including pH and the presence of metal ions^13^. Both *haao-1* knockdown and 3HAA supplementation increased survival of worms exposed to the superoxide generator paraquat (**Fig. 3A**), suggesting that 3HAA is an antioxidant in the context of aging *C. elegans*. We intended to further examine the impact of 3HAA on resistance to exogenous hydrogen peroxide (H_2_O_2_); however, when we added H_2_O_2_ to NGM supplemented with 1 mM 3HAA, we noted a distinct lightening of the red coloration, which we confirmed in solution containing only 3HAA and H_2_O_2_ (**Fig. 3B**). This color change suggested a direct interaction between 3HAA and H_2_O_2_, and we hypothesized that 3HAA may directly degrade hydrogen peroxide. We incubated 3HAA and H_2_O_2_ at a range of concentrations and measured the resulting H_2_O_2_ concentration. 3HAA reduced measurable H_2_O_2_ in a concentration-dependent manner (**Fig. 3C**), and effect that was independent on pH (**Extended Data Fig. 7A**). *C. elegans* release endogenously produced H_2_O_2_ into their environment^14,15^. We found that H_2_O_2_ release increased with age in wild type animals, and that this increase was partially repressed in animals lacking *haao-1* or supplemented with 3HAA (**Fig. 3D**).

**Figure 3.**
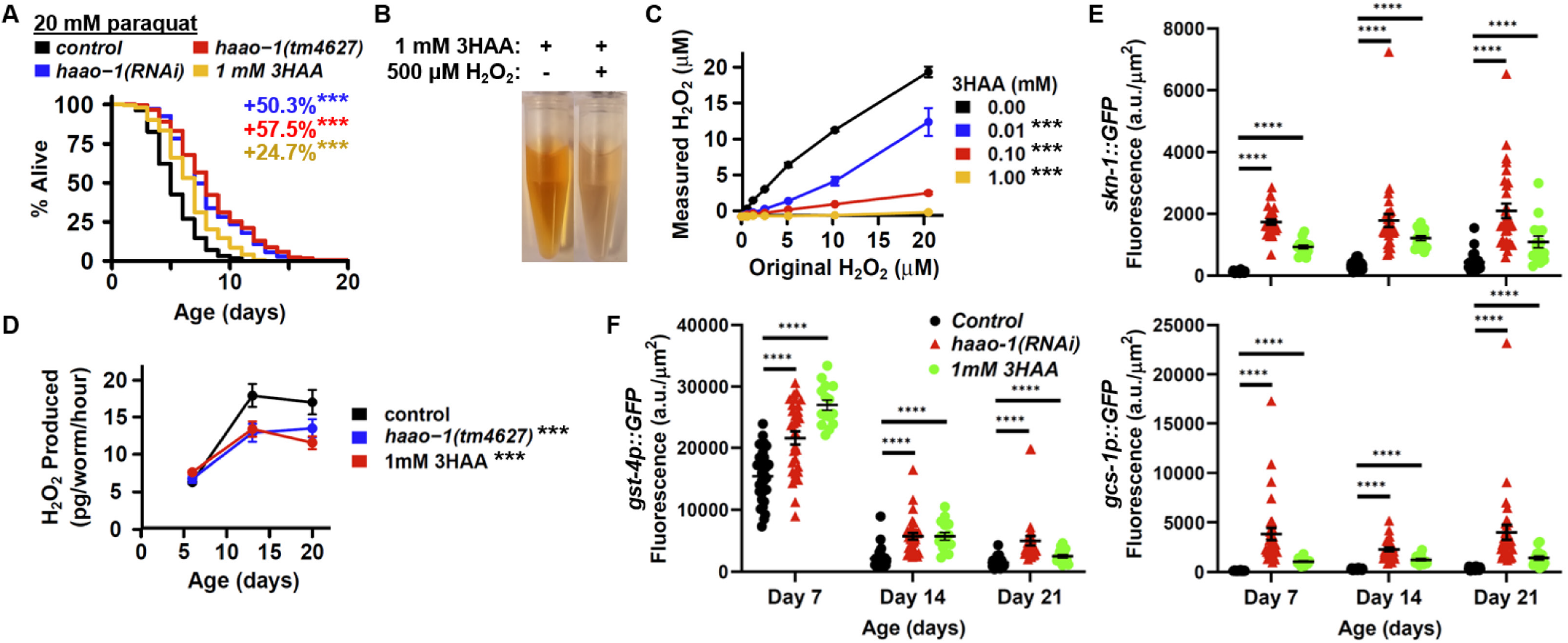
Elevating 3HAA protects against oxidative stress by directly degrading hydrogen peroxide and activating SKN-1-mediated oxidative stress response. (**A**) RNAi knockdown of *haao-1*, deletion of *haao-1 (haao-1(tm4637))*, and media supplemented with 1 mM 3HAA increase survival of *C. elegans* challenged with 20 mM paraquat (colored text indicates change in mean lifespan relative to control). (**B**) 500 μM H_2_O_2_ reduces visible red color of 1 mM 3HAA in water. (**C**) 3HAA reduces detectable H_2_O_2_ in water in a dose-dependent manner. (**D**) Deletion of *haao-1 (haao-1(tm4637))* or 1 mM 3HAA supplementation reduces the age-dependent increase in H_2_O_2_ secretion in *C. elegans*. RNAi knockdown of *haao-1* or 1 mM 3HAA supplementation increases fluorescence in *C. elegans* strains transgenically expressing (**E**) the *skn-1::GFP* fusion protein or (**F**) the *gcs-1p::GFP* or *gst-4p::GFP* promoter activity reporters. All error bars indicate standard error of mean. * p < 0.05, ** p < 0.01, *** p < 0.001.

Our data reveal direct antioxidant properties for 3HAA; however, 3HAA it may also combat oxidative stress by activating endogenous oxidative stress response systems. We examined the impact of 3HAA on four transgenic fluorescent oxidative stress response reporters: (1) green fluorescent protein (GFP) fused to the 3’ end of SKN-1, the *C. elegans* ortholog of the oxidative stress-responsive Nrf2 transcription factor (*skn-1::GFP*), two transcriptional reporters of SKN-1 activity with GFP expression driven by promotors from SKN-1 target genes involved in production of the antioxidant glutathione (*gcs-1p::GFP, gst-4p::GFP*), and a transcriptional reporter of DAF-16 activity with GFP expression driven by the promoter for the antioxidant enzyme superoxide dismutase 3 (*sod-3p::GFP*). Both *haao-1(RNAi)* and 3HAA robustly increased expression of SKN-1::GFP (**Fig. 3E**) and both SKN-1 transcriptional reporters (**Fig. 3F**). *haao-1(RNAi)*, but not 3HAA, slightly activated expression of GFP driven by the SOD-3 promotor (**Extended Data Fig. 7B**), which is consistent with our earlier observation that *haao-1(RNAi)* can extend lifespan in the absence of *daf-16* (**Fig. 1G**). Activation of these oxidative stress reporters by 3HAA was, in some cases, enhanced in worms fed the OP50 strain of *E. coli* relative to the HT115 strain used in RNAi experiments (**Extended Data Fig. 7C**), indicating that kynurenine metabolism may interact with the specific bacteria used as a food source. We concluded that 3HAA has both direct and indirect antioxidant activity *in vivo*.

### Dietary 3HAA supplementation extends healthy lifespan in old mice

In rodents, short-term 3HAA treatment is beneficial in acute models of cardiovascular disease^16^, spinal cord injury^17,18^, asthma^19^, and autoimmune encephalomyelitis^20^. Chronic 3HAA supplementation in the context of normal aging, analogous to our *C. elegans* experiments, has yet to be examined in mammals. We conducted a small study to evaluate the efficacy of 3HAA as therapeutic target to extend healthy lifespan. We fed a small cohort of male C57BL/6J mice a diet supplemented with 0, 312.5, or 3125 ppm 3HAA starting at 27 months of age and measured survival and several metrics of health. Animals on both 3HAA diets were long-lived relative to mice on the control diet, with the 312.5 ppm 3HAA group living the longest (**Fig. 4A**). Low-dose 3HAA significantly improved rotarod performance after 13 weeks, with a trend toward better maintenance of grip strength relative to control mice (**Fig. 4B**). Mice on high-dose 3HAA maintained grip strength over the same 13 weeks better than control mice, with a trend toward improved rotarod performance. Elevated glucose sensitivity is often observed in long-lived populations, such as wild type mice subjected to dietary restriction^21^ or Ames dwarf^22^ mice; however, this association is not universal, as rapamycin can impair glucose sensitivity^23,24^. During glucose tolerance testing conducted 15 weeks after initiating the 3HAA diet, low-dose mice displayed higher glucose both prior to a glucose challenge, and 15 minutes post-challenge, a difference that was not observed during baseline testing (**Extended Data 8A**, **B**). We did not observe notable differences in other parameters measured (**Extended Data Fig. 8C-H**).

**Figure 4.**
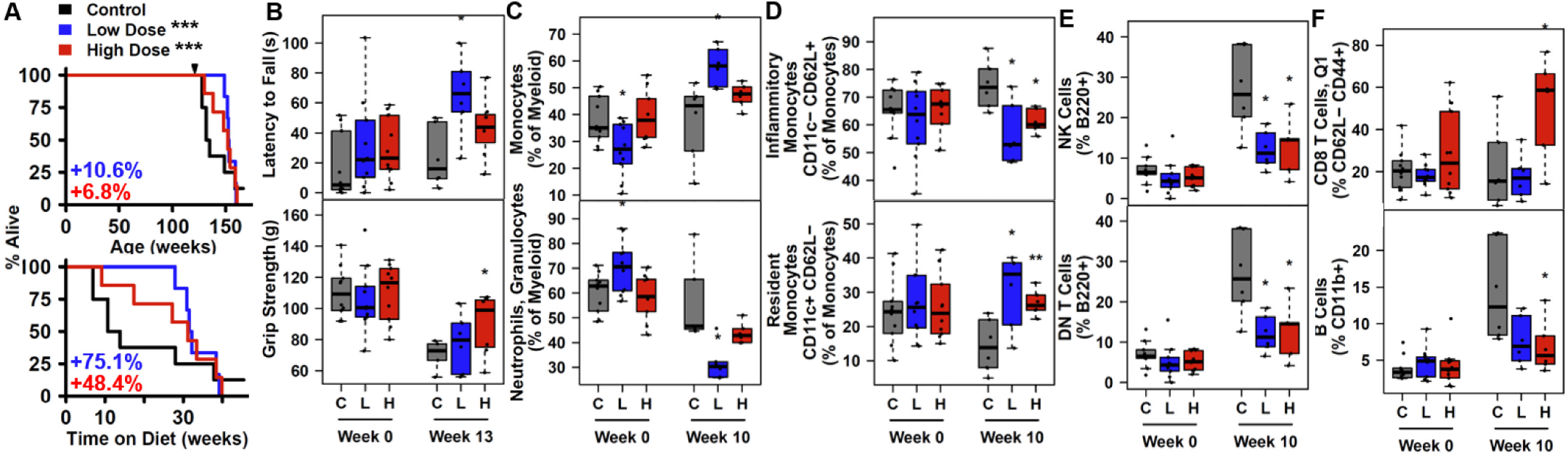
3-hydroxyanthranilic acid (3HAA) extends lifespan, improves health metrics, and promotes a less inflammatory immune cell profile in aging mice. (**A**) C57BL/6J mice fed chow supplemented with 312.5 ppm (low dose) or 3125 ppm (high dose) 3HAA starting at 27 months of age live longer than mice fed control diet. Kaplan-Meier survival curves are displayed as full lifespan (top) or remaining lifespan at the time that 3HAA diet was introduced (bottom). Colored text indicates change in mean lifespan or remaining lifespan relative to control. (**B**) Low dose 3HAA increases rotarod performance, while high dose 3HAA improves grip strength maintenance after 13 weeks. Dietary supplementation with 3HAA alters circulating immune cell profiles after 10 weeks: (**C**) shifting myeloid cells away from neutrophils/granulocytes and toward monocytes, (**D**) shifting monocytes populations away from inflammatory monocytes and toward resident monocytes, (**E**) preventing an increase in both natural killer (NK) and double negative (DN) T cells, and (**F**) increasing CD8 T cell counts while reducing B cell counts (high dose only). * p < 0.05, ** p < 0.01, *** p < 0.001 vs. control diet (Wald test in Cox regression with weighted estimation, panel A; t test, panels B-F).

## Discussion

Here we report the that elevating physiological levels of the kynurenine pathway metabolite 3HAA through genetic inhibition of HAAO or direct 3HAA supplementation extends healthy lifespan in *C. elegans*. The benefits of reduced HAAO activity displayed a complex interaction with established aging pathways, suggesting that the primary mechanism(s) of action are likely orthogonal to the set of pathways examined. We further found that 3HAA increases resistance to oxidative stress by both directly degrading H_2_O_2_ and activating SKN-1-mediated oxidative stress response, providing one molecular mechanism for lifespan extension. Finally, we show that supplementing aging mice with dietary 3HAA is sufficient to promote healthy longevity. Importantly, this last study had a small sample size and was only conducted in male mice, so will require replication; however, our results provide a tentative validation of our invertebrate observations about the benefits of 3HAA during aging in a mammalian system.

The kynurenine pathway is increasingly recognized as an important player in aging and age-associated disease. This study adds to the growing body of work on kynurenines in *C. elegans* aging by implicating *haao-1* as a fourth genetic target capable of influencing longevity that acts via mechanisms that are at least partially distinct from *tdo-2, kynu-1*, and *acsd-1*. Our data support 3HAA as the primary mediator of lifespan extension via *haao-1* knockdown. Studies to date do not exclude the possibility for regulatory feedback between these various kynurenine pathway enzymes and metabolites, which may be important to developing a comprehensive model for the role of kynurenine metabolism in aging. Indeed, our metabolomics data indicate *haao-1* knockdown results in an increase in TRP and KA, but a decrease in anthranilic acid (AA) and xanthurenic acid (XA), suggesting that elevating 3HAA may generate a negative feedback on upstream kynurenine pathway enzymes (**Fig. 1A**, **Extended Data Fig. 1B**). In this same vein, accumulating evidence supports a model in which elevating NAD^+^ is beneficial for healthy aging, including in *C. elegans*^25^. Knockdown of *tdo-2, kynu-1*, or *haao-1* all increase lifespan (**Fig. 1B**) while moderately decreasing NAD^+^ in worms (**Extended Data Fig. 1B**). One implication is that HAAO inhibition and 3HAA supplementation may impart benefits through secondary mechanisms that are distinct, such as 3HAA acting as an NAD^+^ precursor when supplemented in the presence of active HAAO. If this is the case, synergistic benefits may be realized by combining selected kynurenine pathway enzyme inhibitors with strategies to increase physiological NAD^+^ levels, such as supplementing other NAD^+^ precursors or inhibition of NAD^+^-consuming enzymes.

Our data demonstrate that 3HAA is protective against reactive oxygen species in the context of aging *C. elegans*. 3HAA has a complex history with oxidative stress, with earlier studies largely linking 3HAA to ROS generation and/or oxidative damage^26–29^ and more recent studies reporting antioxidant properties for 3HAA^30–37^. These studies vary widely in the specific oxidant properties examined, presence of added metals, cell type, and cellular context (*in vitro* vs. *in vivo)*. 3HAA can auto-oxidize under specific conditions, such as in the presence of copper (Cu^2+^) or iron (Fe^3+^) at alkaline pH^26^. Each of these variables is potentially critical to understanding redox properties of 3HAA in the context of aging cells, and the particular importance of copper and pH were recently confirmed *in silico*^13^. The observations reported here represent the first examination of chronic exposure of whole animals to elevated physiological 3HAA under otherwise standard *C. elegans* culture conditions. Given that 3HAA is protective against oxidative stress and the complex interaction between *haao-1* and genes in the set aging pathways examined (**Fig. 1G**), 3HAA may be protective against other forms of cellular stress.

In summary, this work highlights 3HAA as a novel metabolic target at the intersection of aging, metabolism, and stress response. While earlier work shows that 3HAA can be protective in the context of acute disease in young animals, this is the first demonstration that it can also provide benefits in the context of aging, including imparting health benefits to aging mice, motivating a more detailed evaluation of 3HAA in aging and age-associated disease.

## Acknowledgments

We thank Dr. Aric Rogers, Dr. Jarod Rollins, and Santina Snow at the Mount Desert Island Biological Laboratory (MDIBL) for helpful discussion. We further thank Dr. Rogers for use of the WormLab system. This work was supported by NIH P30AG038070 to Gary Churchill and RK, NIH R35GM133588 to GLS, and the State of Arizona Technology and Research Initiative Fund administered by the Arizona Board of Regents. The mouse study was supported by a pilot award to GLS from The Jackson Laboratory Nathan Shock Center for Excellence in Basic Biology of Aging (NIH P30AG038070). GSL was supported as a JAX Scholar in Aging through The Jackson Laboratory during this work. We gratefully acknowledge the contribution of [optional: name of person] and the [insert Service’s name] Service at The Jackson Laboratory for expert assistance with this publication.

## Materials & Methods

### C. elegans

#### Strains

The following strains were obtained from the *Caenorhabditis* Genetic Center (CGC) at the College of Biological Sciences at the University of Minnesota: *daf-16(mu86)I* (CF1038), *eat-2(ad465)II* (DA465), *hif-1(ia4)V*(ZG31), *rsks-1(ok1255)III*(RB1206), *sir-2.1(ok434)IV*(VC199), *rde-1(ne219) V; kzIs9[(pKK1260) lin-26p::NLS::GFP + (pKK1253) lin-26p::rde-1 + rol-6(su1006)]* (NR222), *lin-15B(n744) X; uIs57[unc-119p::YFP + unc-119p::sid-1 + mec-6p::mec-6]* (TU3335), *rde-1(ne219) V; kbIs7[nhx-2p::rde-1 + rol-6(su1006)]* (VP303), *rde-1(ne300) V; neIs9[myo-3::HA::RDE-1 + rol-6(su1006)]X*(WM118), *wgIs341[skn-1::TY1::EGFP::3xFLAG + unc-119(+)]* (OP341), *dvIs19[(pAF15)gst-4p::GFP::NLS]* (CL2166), *ldIs3[gcs-1p::GFP + rol-6(su1006)]* (LD1171)^38^, *muIs84[(pAD76) sod-3p::GFP + rol-6(su1006)]* (CF1553). Strains *kynu-1(tm4924)X* (FX04627; backcrossed 6x to N2 to create strain GLS129) and *haao-1(tm4627)V* (FX04627; backcrossed 6x to N2 to create strain GLS130) were obtained from the *C. elegans* National Bioresource Project (NBRP) at the School of Medicine at the Tokyo Women’s Medical University. Wild-type (N2) worms were originally obtained from Dr. Matt Kaeberlein (University of Washington, Seattle, WA, USA). Strain RB1206 was generated by the *C. elegans* Gene Knockout Project at the Oklahoma Medical Research Foundation as part of the International *C. elegans* Gene Knockout Consortium^39^. Strain VC199 was generated by the *C. elegans* Reverse Genetics Core Facility at the University of British Columbia as part of the International *C. elegans* Gene Knockout Consortium^39^. Strain OP341 was constructed as part of the Regulatory Element Project, part of modENCODE^40^. Strain *haao-1(tm4627) Vkynu-1(tm4924) X*(GLS147) was generated by crossing GLS129 to GLS130. Strains *haao-1(syb2665)::wrmscarlet* (PHX2665) and *kynu-1(syb2691)::wrmscarlet* (PHX2691) were created using CRISPR/Cas9 precise gene insertion by SunyBiotech Corporation.

#### Media and culture

We maintained worms on solid nematode growth media (NGM) seeded with *Escherichia coli* bacteria at 20°C as previously described^41^ except where otherwise noted. RNAi experiments used *E. coli* strain HT115, while all other experiments used strain OP50. We conducted RNAi feeding, lifespan, healthspan, brood size, paralysis, and aggregate quantification assays according to standard protocols. All worms were transferred to NGM plates containing 50 μM 5-fluorodeoxyuridine (FUdR) starting at the L4 larval stage to prevent reproduction.

#### RNA interference (RNAi)

All experiments were conducted on NGM containing 1 mM Isopropyl β-D-1-thiogalactopyranoside (IPTG) to activate production of RNAi transcripts and 25 μg/mL carbenicillin to select RNAi plasmids and seeded with live *E. coli* (HT115) containing RNAi feeding plasmids. Worms were age-synchronized via timed egg laying (with the exception of oxidative stress assays, which were synchronized via bleach prep) at the experimental temperature and transferred to plates containing 50 μM 5-fluorodeoxyuridine (FUdR) to prevent reproduction at the L4 larval stage as previously described^41^.

#### 3HAA supplementation

3HAA supplementation was achieved by adding the appropriate mass of solid 97% pure 3HAA (Sigma-Aldrich, catalog number 148776) to NGM media during preparation prior to autoclaving.

#### Lifespan analysis

Lifespan and paralysis experiments were conducted as previously described^41^. Briefly, adult animals were maintained on NGM RNAi plates with FUdR throughout life. Each animal was examined every 1-2 days (25°) or 2-3 days (15°C, 20°) by nose- and tail-prodding with a platinum wire pick. Animals were scored as dead if they failed to react to prodding. Live and dead animals were counted, and dead animals removed from the plate at each examination. Animals displaying vulva rupture were included in all analyses, while worms that left the surface of the plate were excluded.

Each lifespan experiment included test groups consisting of 105-150 worms (3 plates with 35-50 worms/plate). Each experiment included a negative control test group fed *E. coli* strain matched to the test conditions. For lifespan epistasis and tissue-specific RNAi experiments, each combination of candidate gene RNAi and mutant/transgenic strain was measured in three independent experiments, each including: wild type worms fed *EV(RNAi)*, wild-type worms fed candidate RNAi, mutant worms fed *EV(RNAi)*, and mutant worms fed candidate RNAi. P-values for statistical comparison of lifespan between test groups were calculated using the log-rank test (*survdiff* function in the R “survival” package) with Holm multiple test correction applied to comparisons made within each experiment.

#### Kynurenine pathway metabolite quantification

##### Metabolite extraction

Worms were grown plates containing live *E. coli* (HT115) containing RNAi feeding plasmids targeting *kynu-1(RNAi), haao-1(RNAi)*, or *tdo-2(RNAi)* at 15°C under conditions identical to those used in the lifespan studies. On day 4 of adulthood, ~100 worms/sample were collected from the plates and washed twice with M9 buffer (21.6 mM Na_2_HPO_4_ 22mM KH_2_PO_4_ 85.6 mM NaCl, 1mM MgSO_4_). Excess M9 was removed and the worm pellets flash frozen in liquid nitrogen and stored at −80°C prior to metabolite extraction. To extract metabolites, 1 ml of extraction buffer (2:2:1 AcN:MeOH:H2O + internal standards [0.2 ng/ul 1-Napthylamine + 0.2 ng/ul 9-anthracene carboxylic acid]) was added to each sample. Samples were homogenized mechanically and sonicated to dissolve worm pellet and placed at −20°C overnight to precipitate metabolites. Samples were then centrifuged at 20,000 x g for 15 min to pellet protein, and the supernatant transferred to a new tube and dried without heat. Metabolites were resuspended in a solution of 10% acetonitrile in water, both Optima^®^ grade for mass spectrometry analysis.

##### Relative quantification via mass spectrometry (MS)

Samples were run on an Agilent 6530 Q-TOF equipped with a microflow liquid chromatography (LC) system. Separation by reverse-phase LC (RPLC) with a C18 column (Agilent Poroshell, 2.1mm x 50mm) over a 20 minute gradient resulted in separate peaks for each analyte. Detection was in positive ion mode over a mass range of 200-1700 m/z with internal reference ions. Collision-induced dissociation (CID) was employed to confirm the identification of each analyte by mass and retention time. For preliminary analysis, both automated detection and fragmentation of all analytes, as well as detection and fragmentation of targeted analytes were employed. Both resulted in sufficient fragmentation for identification and sufficient intensity for quantification. Subsequent analysis employed quantitative scans for confirmation of relative abundance.

##### Data analysis

Samples were extracted for target ions using ACD Labs MSWorkbookSuite IntelliTarget feature (http://www.acdlabs.com/products/spectrus/workbooks/ms/msworkbooksuite/). Briefly, a database for the targets of interest was generated and populated using standard compounds as well as library reference data. Targets were extracted and quantified with normalization against an internal standard. Scans were verified manually at both the first and second stages of MS (MS1 and MS2).

We gratefully acknowledge the contribution of Dorothy Ahlf Wheatcraft in the Protein Sciences Service at The Jackson Laboratory for expert assistance with this portion of the work.

#### Brood size

To measure brood size, 10 worms at the L4 larval stage were placed on individual NGM RNAi plates lacking FUdR and allowed to lay eggs. Worms were transferred to a new plate every 12 hours at 25°C or every 24 hours at 15°C until worms ceased laying eggs. The number of progeny was counted on each plate following a 2 day incubation to allow eggs to hatch. Student’s t-tests were used to determine significance in observed differences between target and EV RNAi at each age, and between total progeny counts.

#### Healthspan

Healthspan data was collected in three independent experiments using the WormLab system (Version 3.1, MBF Bioscience, Williston, VT) at 7, 14, 21, and 25 days of age (15°C) and 4, 8, 11, and 15 days of age (25°C). For speed assays, videos were captured for worms directly on NGM experiment plates. Worms were motivated to begin moving by dropping the plate onto the WormLab video capture stage from a height of ~0.5 inches. Video capture was started immediately and allowed to run for 1 minute, 15 seconds. Worms reacted to the plate drop by moving rapidly for approximately 20 seconds then returning to a normal foraging behavior. For “motivated speed”, worms were tracked using the WormLab software for the first 15 seconds of each video. Worms present for at least 14 seconds were selected for analysis. For “unmotivated speed”, worms were tracked between 30 seconds and 1 minute, 15 seconds. Worms present for at least 30 seconds were selected for analysis. For thrashing, worms were suspended in a droplet of M9 buffer on unseeded NGM plates and video captured for 45 seconds. Worms present for at least 15 seconds were selected for analysis. Average speed, thrash frequency, body length, and body width for each worm was exported from the WormLab software to R for analysis. The WormLab software tended to misidentify dark sections of the plate as worms. To filter these errors, we initially removed any identified object that fell outside of 1.2 standard deviations in worm body length, width, or area. Based on manual inspection, this threshold removed non-worm objects and poorly identified worms without removing correctly identified worms. Student’s t-tests were used to determine significance in differences in each phenotype between target and EV RNAi at each age.

#### Red pigmentation imaging

Images of wild type and *haao-1(tm4627) C. elegans* to illustrate accumulation of red pigment were taken using a Samsung S9+ Smartphone using stock camera software operating in professional mode attached to a Zeiss Stemi 508 stereo dissection microscope using a 3d-printed cell phone mount. All camera settings were constant across images.

#### Paraquat toxicity assay

To assess paraquat toxicity, survival assays were conducted as described above with media containing 20 mM paraquat. All animals were scored daily for lifespan.

#### Hydrogen peroxide (H_2_O_2_) measurement

H_2_O_2_ was quantified using Invitrogen™ Amplex™ UltraRed Reagent (ThermoFisher Scientific catalog number A36006) according to manufacturer instructions, including use of the optional Invitrogen™ Amplex™ Red/UltraRed Stop Reagent (ThermoFisher Scientific catalog number A33855). For *in vitro* H_2_O_2_ degradation assays, stock solutions of 3HAA and H_2_O_2_ in water were mixed to achieve the indicated concentrations and incubated at room temperature for 30 minutes prior to quantification. For *in vivo* H_2_O_2_ in *C. elegans*, age-synchronized populations were grown as described above. 100 worms/sample were collected on day 6, 13, and 20 from egg with 5 technical replicates per test condition. Worms were collected and washed 2 times with M9 0.05% Triton X-100, and incubated at 20°C with shaking in 1x reaction buffer (50mM NaH2Po4, 0.05% Triton X-100 pH 7.4) for 3 hours. After incubation, the worms were allowed to settle to the bottom of the tube (3-5 minutes), after which the supernatant was collected for quantification.

#### Fluorescence oxidative stress reporter assay

Age-synchronized populations of *C. elegans* were maintained as described above. At the designated ages, 1-20 animals were manually transferred to a drop of 25 mM sodium azide (NaN_3_) on slides prepared with 6% agarose pads and imaged using a Leica M205 FCA Fluorescent Stereo Microscope equipped with a Leica K6 sCMOS monochrome camera. Identical imaging settings were maintained across timepoints for each strain. Each experiment was repeated in biological triplicate.

### Mice

#### Lifespan study

Male C57BL/6J mice (stock # 000664) were obtained from The Jackson Laboratory and aged to 27 months while being housed in a pathogen-free room with a 12:12 hour light:dark cycle and given a standard rodent diet (LabDiet 5KOG) and acidified water. At 27 months, animals were randomly assigned (n=10 per group) to either a control diet (LabDiet 5LG6-JL), a diet with 312.5 ppm 3HAA (low dose; Sigma-Aldrich, catalog number 148776), or a diet with 3,125 ppm 3HAA (high dose). In addition, blood and urine were collected at both time points to for blood chemistry, urine chemistry, and to determine blood cell composition using flow cytometry (following a standardized protocol for mouse aging studies that is previously described^42^). We gratefully acknowledge the contribution of Will Schott in the Flow Cytometry Service at The Jackson Laboratory for expert assistance with this portion of the work. Mice were kept until their natural death or until humanely euthanized because they reached a moribund state. All animal experiments were performed in accordance with the National Institutes of Health Guide for the Care and Use of Laboratory Animals (National Research Council) and were approved by The Jackson Laboratory’s Animal Care and Use Committee.

##### Statistics

To evaluate statistical significance in mouse lifespan, we first constructed a Cox proportional hazard regression model (*coxph* function in R “survival” package); however, this model failed quality control because it did not meet the proportional hazard assumption (p < 0.05 using *cox.zph* function in the R “survival”), apparently driven by a single long-lived outlier in the control mouse group (*ggcoxdiagnostics* function in the R “survminer” package). To address this issue, we instead employed weighted estimate in Cox regression (*coxphw* function in the R “coxphw” package), a method robust to violations of the proportional hazard assumption^43^. P-values reflect the results of a Wald test performed by the *coxphw* function on pair-wise comparisons of diet groups using this approach.

#### Health and behavioral assessment

Grip strength and latency of fall from a rotarod were measured at both at baseline (27 months) and 30 months as previously described^42^ (Ackert-Bicknell, 2015). Deficit accumulation frailty index (FI) is a metric of general health and resilience that incorporates assessment of 30 physical traits into a single index. FI was measured for all surviving mice at baseline (27 months) and every 2 weeks until a final assessment at 30 months as previously described^44^. Body weight measurement was collected as part of FI assessment. The Y-maze assay to quantify number of arm entries and spontaneous alternations was performed as previously described^44^. We gratefully acknowledge the use of facilities and equipment through the Center for Biometric Analysis at The Jackson Laboratory in this portion of the work.

#### Glucose tolerance test

Food access was removed for all mice 4 hours prior to glucose tolerance test. Glucose, 2 mg/g body weight (G8769; Sigma-Aldrich, St. Louis, MO) was injected into the intraperitoneal cavity. Blood glucose was measured immediately prior to glucose injection (0 minutes) and 15, 30, 60, 120, and 180 minutes post-injection from blood from the tail vein using a NovaStatStrip Xpress handheld glucometer (Nova Biomedical, Waltham, MA).

#### Clinical chemistry

Triglycerides, cholesterol, and blood urea nitrogen (BUN) concentrations were measured in serum separated from whole blood using a Beckman AU680 Chemistry Analyzer, as were urine albumin and creatinine concentrations. Actual mouse albumin concentrations were calculated by linear regression from a standard curve generated with mouse albumin standards (Kamiya Biomedical Company, Seattle, WA)^42^. We gratefully acknowledge the contribution of Todd Hoffert in the Histopathology Service at The Jackson Laboratory for expert assistance with this portion of the work.

#### Body composition

Whole body composition (fat and lean tissues) was measured in live, conscious mice by nuclear magnetic resonance using an Echo MRI Analyzer (Echo Medical Systems, Houston, TX).

