## Extended Data Figures for "3-hydroxyanthranilic acid – a new metabolite for healthy lifespan extension"

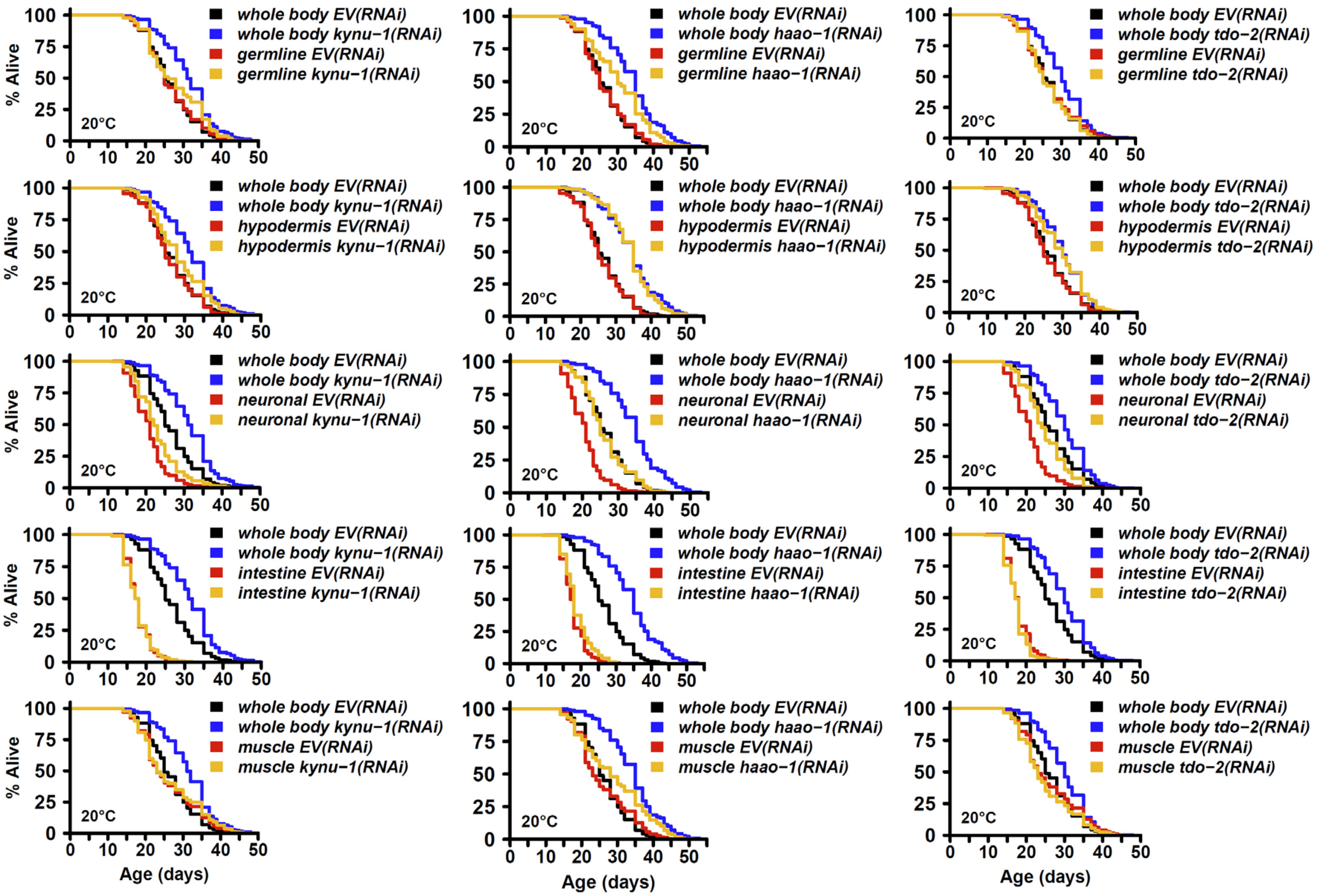


**Extended Data Figure 2. The impact of *kynu-1*, *haao-1*, and *tdo-2* knockdown on lifespan show distinct patterns of tissue dependence.** RNAi knockdown *tdo-2*, *kynu-1*, or *haao-1* in wild type animals compared to mutants or transgenic animals modified such that RNAi is only effective in specific tisues: germline (strain MAH23), hypodermis (strain NR222), neurons (strain TU3335), intestine (strain VP303), or muscle (strain WM118). Each panel shows the impact on *C. elegans* survival for the indicated kynurenine pathway RNAi vs. *EV(RNAi)* in wild type (N2) animals and one strain with active RNAi only in the indicated tissue.

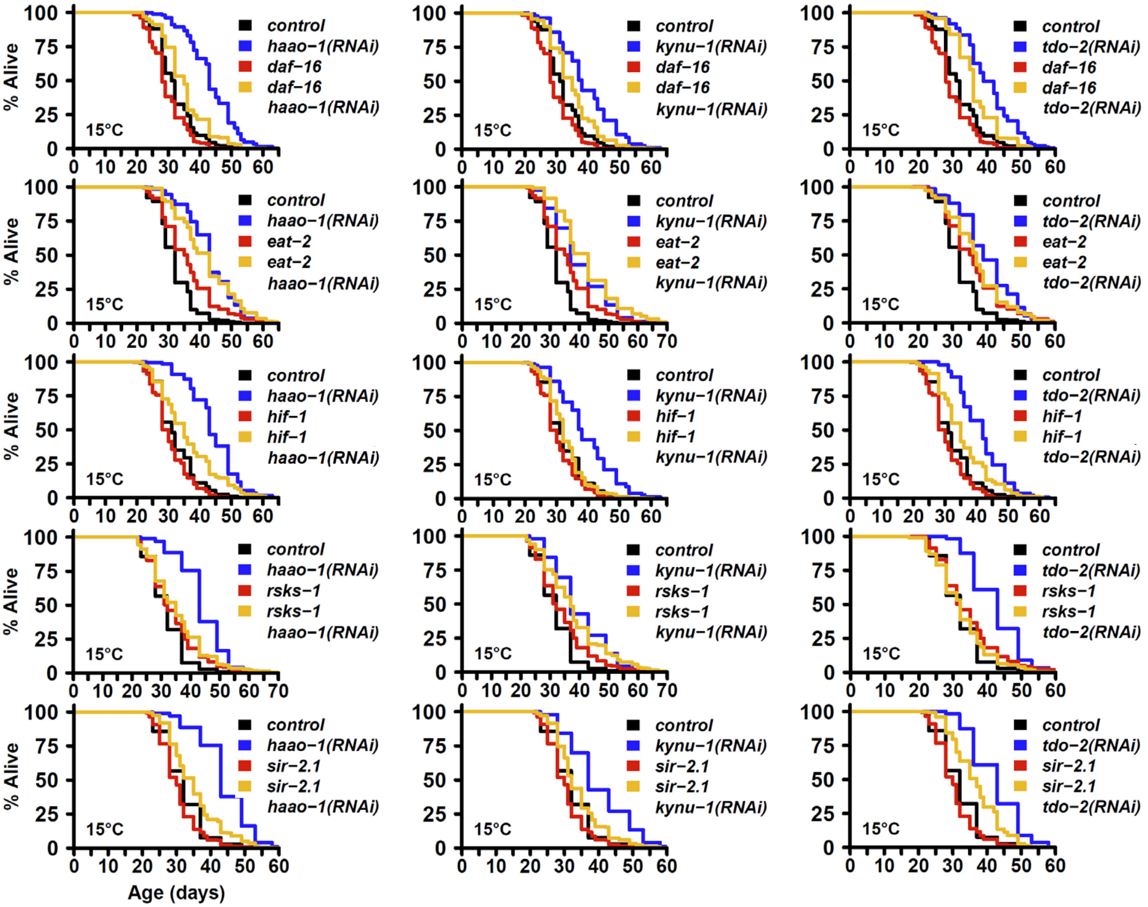


**Extended Data Figure 3. *kynu-1*, *haao-1*, and *tdo-2* display distinct interaction patterns with established aging pathways at 15°C.** Genetic interaction between *tdo-2*, *kynu-1*, and *haao-1* with established aging pathways. Each panel shows the impact on lifespan of RNAi knockdown of either *kynu-1*, *haao-1*, or *tdo-2* vs. *EV(RNAi)* in both wild type (N2) worms and worms with loss-of-function mutations in the following genes with a previous link to aging: *daf-16(mu86)* (strain CF1038), *eat-2(ad465)* (strain DA465), *hif-1(ia4)* (strain ZG31), *rsks-1(ok1255)* (strain RB1206), *sir-2.1(ok434)* (strain VC199).

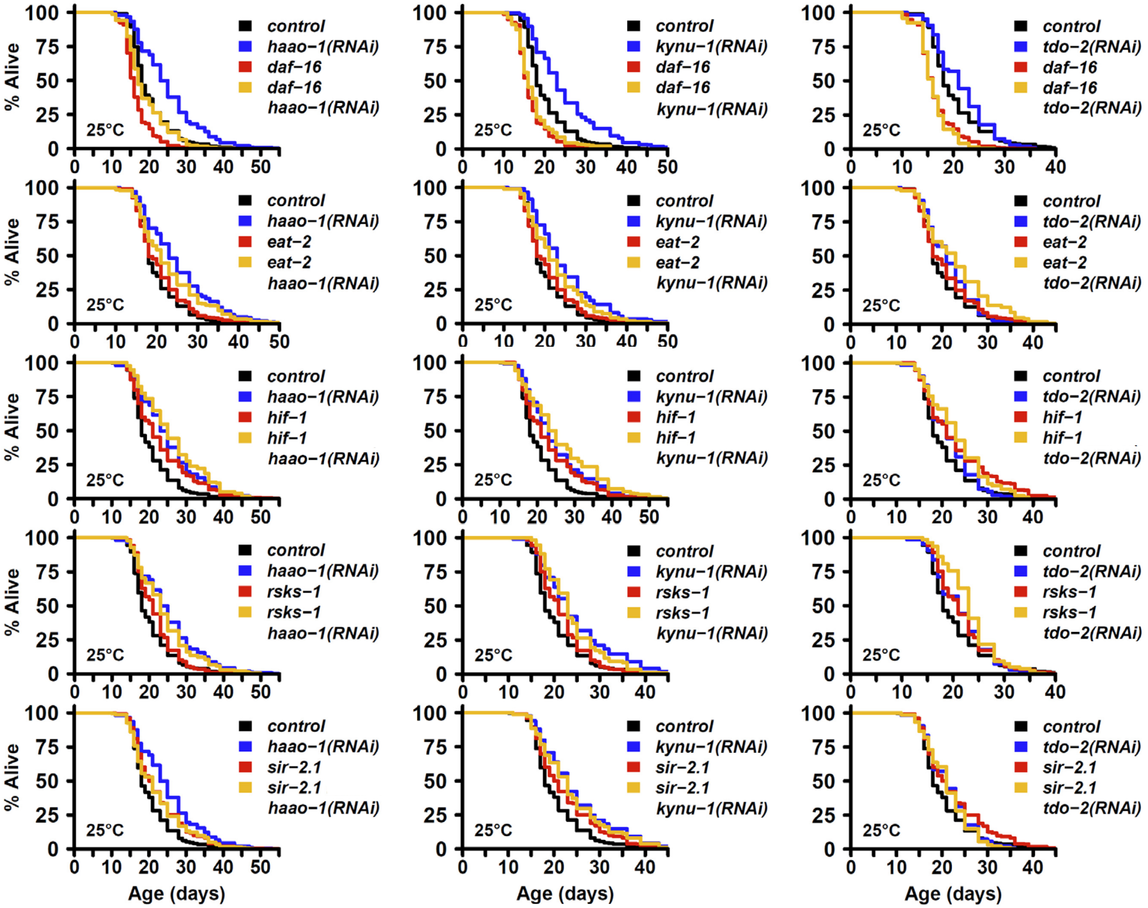


**Extended Data Figure 4. *kynu-1*, *haao-1*, and *tdo-2* display distinct interaction patterns with established aging pathways at 25°C.** Genetic interaction between *tdo-2*, *kynu-1*, and *haao-1* with established aging pathways. Each panel shows the impact on lifespan of RNAi knockdown of either *kynu-1*, *haao-1*, or *tdo-2* vs. *EV(RNAi)* in both wild type (N2) worms and worms with loss-of-function mutations in the following genes with a previous link to aging: *daf-16(mu86)* (strain CF1038), *eat-2(ad465)* (strain DA465), *hif-1(ia4)* (strain ZG31), *rsks-1(ok1255)* (strain RB1206), *sir-2.1(ok434)* (strain VC199).

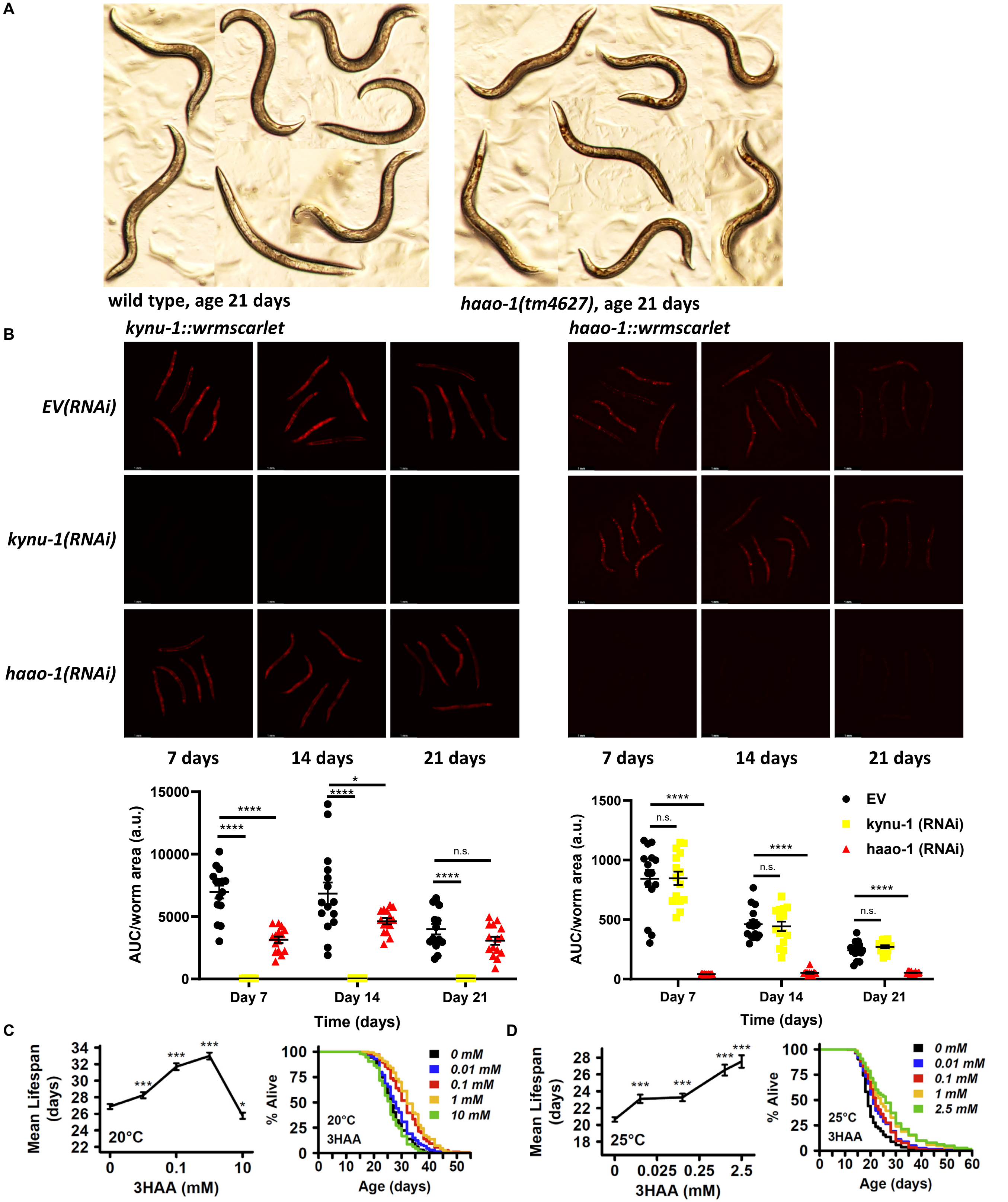


**Extended Data Figure 5. 3HAA is upregulated in animals with reduced *haao-1* and is sufficient to extend lifespan.** (**A**) *haao-1(tm4627* (GLS130; right) *C. elegans* accumulate a red coloration throughout their body as they age that is not present in wild type (N2; left) animals (brightfield images; age 21 days). (**B**) *kynu-1(RNAi) and haao-1(RNAi)* efficiently knockdown expression of transgenic fluorescent fusion reports *kynu-1::wrmscarlet* and *haao-1::wrmscarlet*, respectively. Top panels show representative images of the indicated worm (5 worms/panel). Charts show quantified fluorescence intensity for individual animals in each group. (**C**,**D**) Mean lifespan dose response curves (3HAA) and Kaplan-Meier survival curves (right) for *C. elegans* grown on media supplemented with 3HAA at (**C**) 20°C and (**D**) 25°C. * $p < 0.05$, ** $p < 0.01$, *** $p < 0.001$ vs. *EV(RNAi)* (t test, panel B) or 0mM 3HAA (log-rank test; panels C,D).

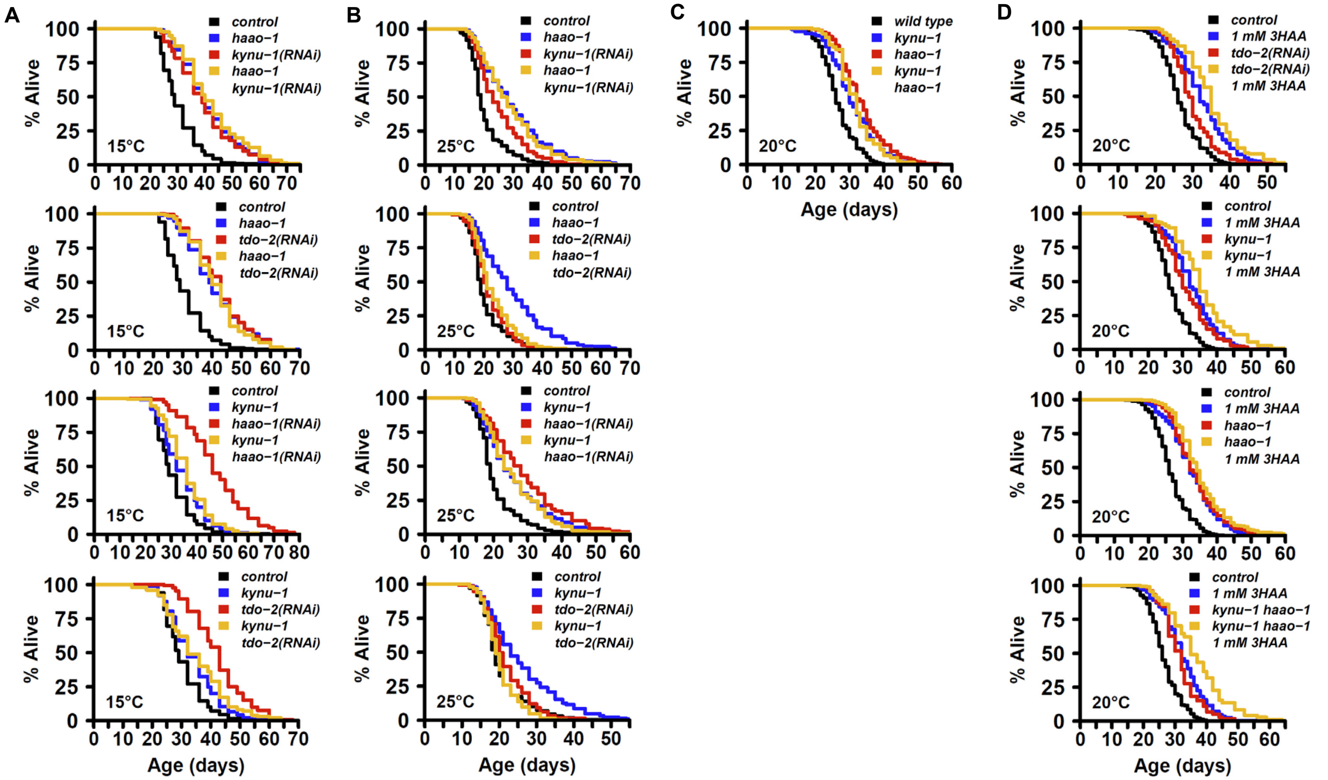


**Extended Data Figure 6. Interaction between kynurenine pathway genes and 3HAA.** Genetic interaction between *tdo-2*, *kynu-1*, and *haao-1* with respect to lifespan at (**A**) 15°C, (**B**) 25°C, and (**C**) 20°C. (**D**) 1mM 3HAA extends lifespan to a degree similar to wild type *C. elegans* in animals deficient for *tdo-2* (top left) and *kynu-1* (bottom left), but not *haao-1* (top right). Knockout of *kynu-1* rescues the ability of 3HAA to extend lifespan in *haao-1* deficient animals (bottom right). In all panels, *haao-1* refers to the *haao-1(tm4627)* knockout strain and *kynu-1* refers to the *kynu-1(4924)* knockout strain, while *tdo-2(RNAi)*, *kynu-1(RNAi)*, and *haao-1(RNAi)* refer to knockdown of the respective genes by RNAi.

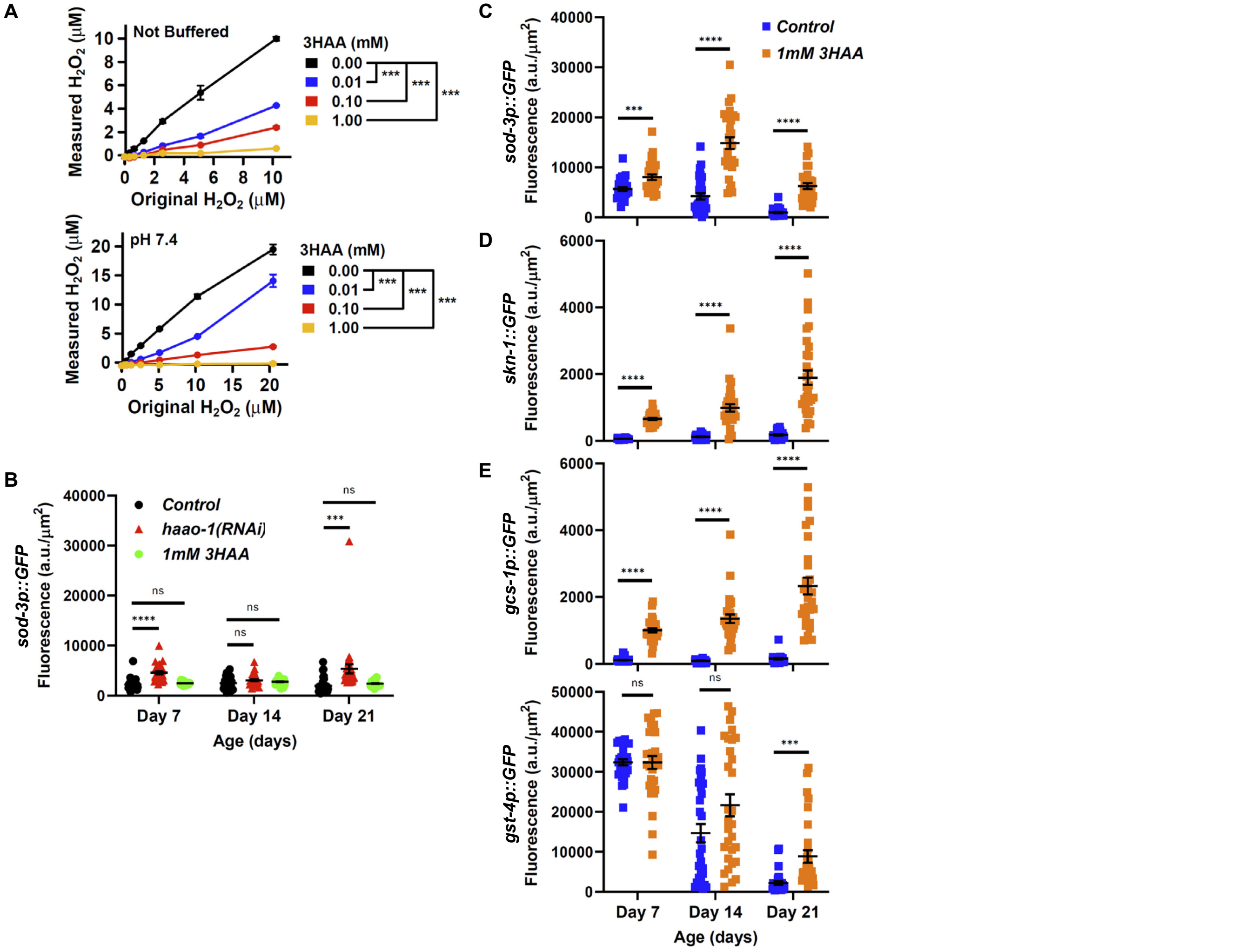


**Extended Data Figure 7. 3HAA degrades hydrogen peroxide and activates oxidative stress response pathway in *C. elegans*.** (**A**) 3HAA reduces detectable $H_2O_2$ in a dose-dependent manner in both unbuffered (top) or buffered (bottom; pH 7.4) solutions. (**B**) RNAi knockdown of *haao-1,* but not 1 mM 3HAA supplementation, mildly increases expression of a transgenically-expressed fluorescent DAF-16 transcriptional reporter (*sod-3p::GFP*) in worms fed the HT115 strain of *E. coli*. (**C**) 3HAA supplementation does activate expression on the OP50 strain of *E. coli*. 3HAA supplementation increases expression of (**D**) *skn-1::GFP* fusion protein and (**E**) the *gcs-1p::GFP* or *gst-4p::GFP* promoter activity reporters in transgenic worms fed *E. coli* strain OP50. All error bars indicate standard error of mean. * $p < 0.05$, ** $p < 0.01$, *** $p < 0.001$ vs. 0 mM 3HAA (ANOVA; panel A) or age-matched control (t test; panels B-E).

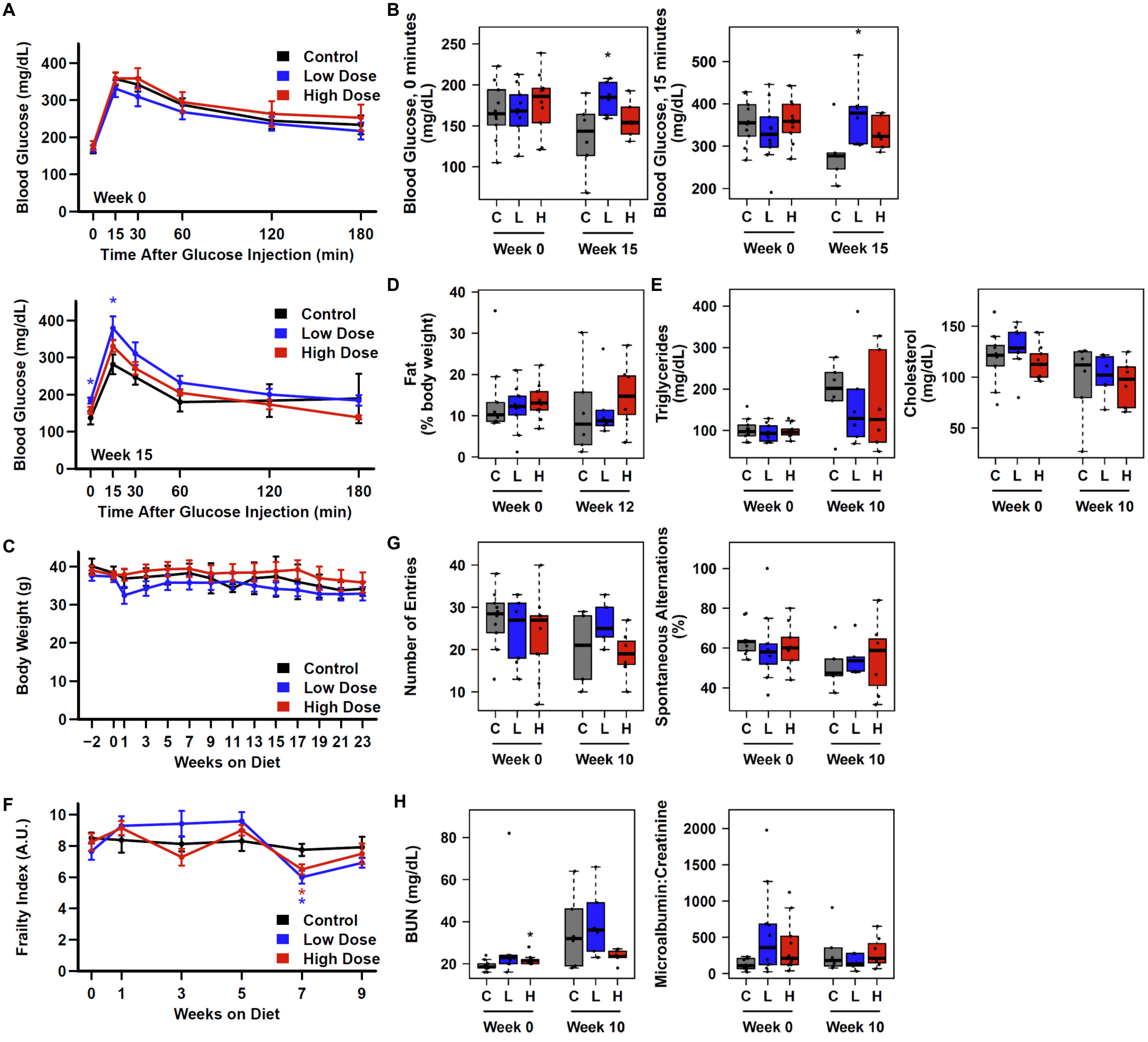


**Extended Data Figure 8. Additional detail on dietary 3HAA on metabolic parameters in aged mice.** (**A,B**) 27-month-old C57BL/6J mice fed chow supplemented with 312.5 ppm (low dose) but not 3125 ppm (high dose) 3HAA for 15 weeks display elevated fasting glucose and 15-minute glucose response. 3HAA supplemented diet does not substantially alter (**C**) body weight after 23 weeks, or (**D**) body composition, or (**E**) circulating triglycerides, cholesterol, (**G**) Y-maze performance, (**F**) frailty index, or (**H**) blood urea nitrogen (BUN) by week 10. * $p < 0.05$, ** $p < 0.01$, *** $p < 0.001$ vs. time-matched control diet (t test).
